# Two-target quantitative PCR to predict library composition for shallow shotgun sequencing

**DOI:** 10.1101/2020.09.21.304006

**Authors:** Matthew Y. Cho, Marc Oliva, Anna Spreafico, Bo Chen, Xu Wei, Yoojin Choi, Rupert Kaul, Lillian L. Siu, Bryan Coburn, Pierre H. H. Schneeberger

## Abstract

Shotgun sequencing enables retrieving high resolution information 40 from complex microbial communities. However, the technique is limited by missing information about host-to-microbe ratios observed in different sample types. This makes it challenging to plan sequencing experiments, especially in the context of high sample multiplexing and/or limited sequencing output. We evaluated a qPCR-based assay to predict host-to microbe ratio prior to sequencing. Using a two-target assay aimed at conserved human and bacterial genes, we predicted human-to-microbe ratios in two sample types and validated it on independently collected samples. The assay enabled accurate prediction for a broad range of sample compositions.

## Introduction

Shotgun sequencing allows interrogation of the metagenomic composition of ecological niches and has been increasingly utilized to characterize human-associated microbial communities. Shallow shotgun sequencing – sequencing to a per-sample read depth of 10^5^ to 10^6^ reads – provides taxonomic resolution greater than 16S amplicon sequencing and functional characterization of metagenomes, while being less expensive than whole genome sequencing or deep sequencing (typically 10^7^ to 10^9^ reads/sample) (1). However, there is a trade-off between cost and adequacy which is especially problematic for samples of variable ratios of host to microbial DNA, where microbial reads may be displaced by human reads in a mixed sample (2). While this is generally not a concern for samples with high bacterial load, such as stool samples, samples with low or variable microbial DNA relative to human DNA are common in other regions of the body, such as the lung, nasopharynx, stomach, and duodenum (2, 3, 4, 5). Microbial taxonomic and functional analyses of metagenomic data require sufficient reads to draw robust conclusions. The ability to predict the proportion of microbial reads prior to sequencing would allow researchers to customize sequencing strategies for desired analyses, while optimizing the cost and time spent on metagenomic sequencing.

In this study, we used quantitative PCR to predict the ratio of human to microbial reads obtained from sequencing using three targets: the 16S rRNA gene, 18S rRNA gene, and human beta-actin (ACTB) to quantitate DNA of bacterial, fungal, or human origin, respectively (6-8). We compared the ratios of bacterial to human DNA determined via qPCR to the percent microbial/human DNA determined via shallow shotgun sequencing in samples with variable bacterial DNA. We derived a prediction model from oropharyngeal swabs and stool samples, and evaluated it in a set of independently collected samples, including rectal swabs and vaginal secretion samples. Finally, we generated an easy-to-use tool based on qPCR data to predict sample composition and sequencing depth required given a desired analytical outcome.

## Results and Discussion

To assess the impact of shallowing sequencing depth on different bacterial DNA proportions, we rarefied shotgun sequencing data from 4 sample types – stool, oropharyngeal, rectal, and vaginal – to depths of 1000 to 1 million reads/sample. We then determined the alpha diversity of each rarefaction using three metrics: richness, Shannon index, and Berger-Parker index. Alpha diversity decreased in a sample type-specific manner as sequencing depth decreased (**Fig. 1**). Notably, while vaginal samples have the lowest alpha diversity in all three metrics of the four sample types, alpha diversity decreased at the slowest rate as sequencing depth decreased (**Fig. 1**). Conversely, while rectal swab samples had similar Shannon index and Berger-Parker index values at 10^6^ microbial reads to oropharyngeal and stool samples, alpha diversity in rectal samples diminished at a greater rate as sequencing depth decreased (**Fig. 1B-C**). Since this effect is sample type-specific, it is critical to predict sample composition *a priori* to ensure sufficient reads for the desired analysis for the given sample type.

**Figure 1.**
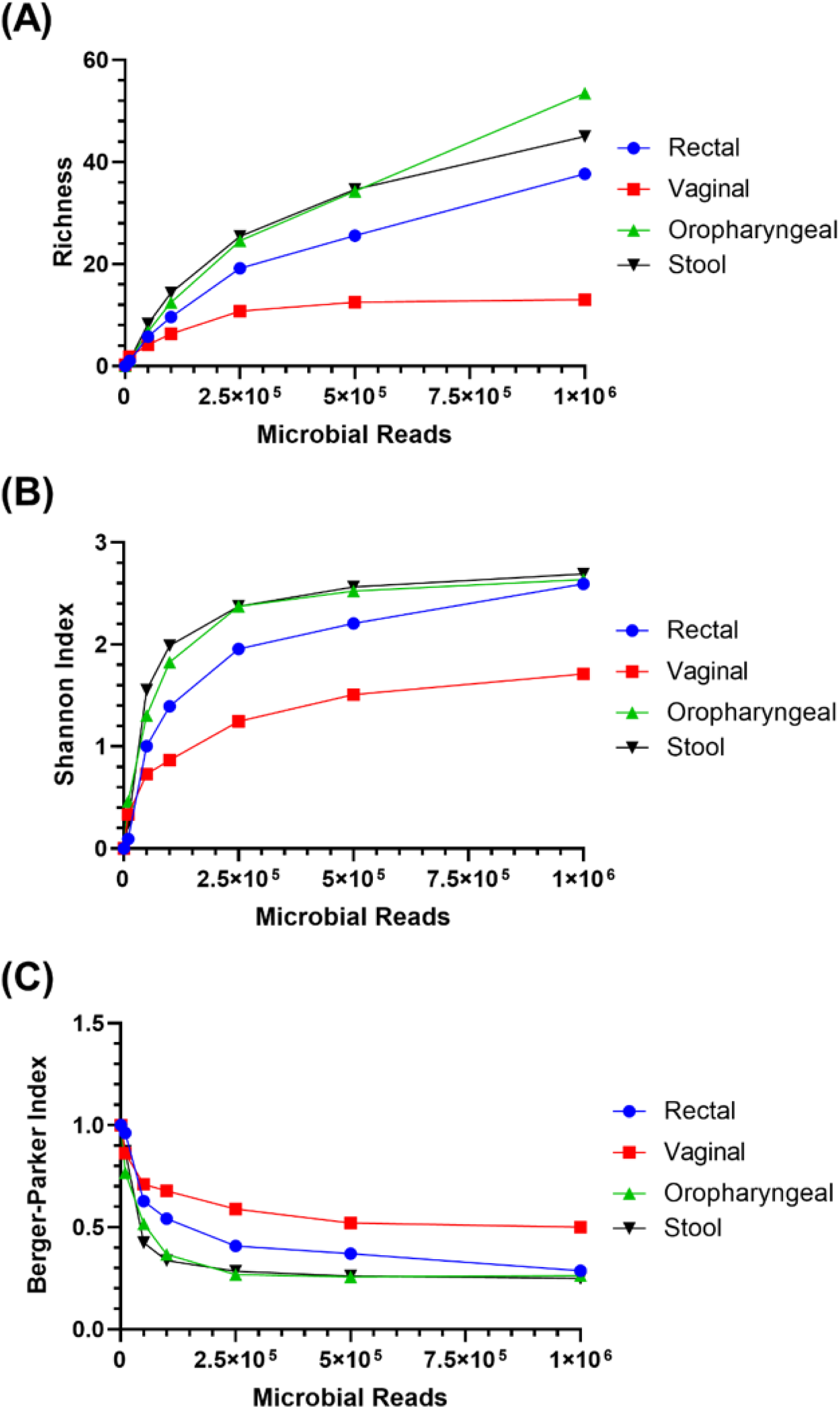
Alpha diversity indices are shown across a range of simulated sequencing depths from 1E3 to 1E6 reads per sample. (A) Sample-specific rarefaction curves of species richness. (B) Shannon index calculated across a range of rarefactions, by sample type. (C) Sample dominance, measured with the Berger-Parker index, across a range of sequencing depths, stratified by sample type.

qPCR is a widespread and robust technique available in many molecular biology laboratories. Its availability as well as cheap associated costs, especially compared to experiments involving high-throughput sequencing techniques, makes it an ideal candidate to use to predict sample composition prior to sequencing. In this study, we assessed the potential of qPCR to predict sample-specific ratios of human to microbe DNA using different amplification targets. Using a multivariate approach, 5 models were generated mapping 16S rRNA gene, 18S rRNA gene, and human beta-actin (ACTB) qPCR-derived cycle thresholds (Ct) to observed percentage of microbial reads for a sample set consisting of oropharyngeal swabs and stool samples. Microbial reads were defined as any read which did not align/match with a human genome reference. The following models were tested: (A) a linear fit using 16S rRNA gene and ACTB Ct values, (B) a linear fit using 16S rRNA gene, 18S rRNA gene, and ACTB Ct values, (C) a linear fit using logit transformed 16S rRNA gene and ACTB Ct values, (D) a linear fit using logit transformed 16S rRNA gene, 18S rRNA gene, and ACTB Ct values, and (E) a nonlinear regression model based on the logistic growth equation using 16S rRNA gene and ACTB Ct values (**Supplementary figure 1A**). We compared goodness-of-fit for each model and observed R^2^-values of 0.880, 0.880, 0.920, 0.920, and 0.990 for models A – E, respectively (**Figure 2A, Supplementary figure 1A**). Observed residuals had a min-max range of 67.56, 68.50, 58.93, 59.07, and 42.61 for models A – E, respectively (**Supplementary figure A**). Based on these findings, model E turned out to be the best fitting model to predict sample composition using qPCR, with an equation of % microbial reads = (2.7201549)/((99.50267)*e^(−0.7218*(ACTB-16S))+ 0.02733). In addition, 18S rRNA Ct value was not found to be an informative predictor and was hence removed from the model. In **Figure 2B**, we show the goodness-of-fit and residuals observed with model E across the range of qPCR differences (−8.16% to +34.45%). We observed homogeneous fit and variance indicating that the model performs well for all observed host to microbe DNA ratios. However, we also observed that the model loses accuracy at each end of the range due to the s initial dataset used and sigmoidal curve generated, with limits approximately at 4% and 98%. This bias is likely introduced at different steps of the process. For instance, sequencing error, and resulting false negative and positive hits when mapping reads to the human database are likely to account for this bias. Another potential source of bias could be introduced by the carryover of contaminants between sequencing runs, hence resulting in a composition change which is not picked up by the qPCR conducted *a priori*.

**Figure 2.**
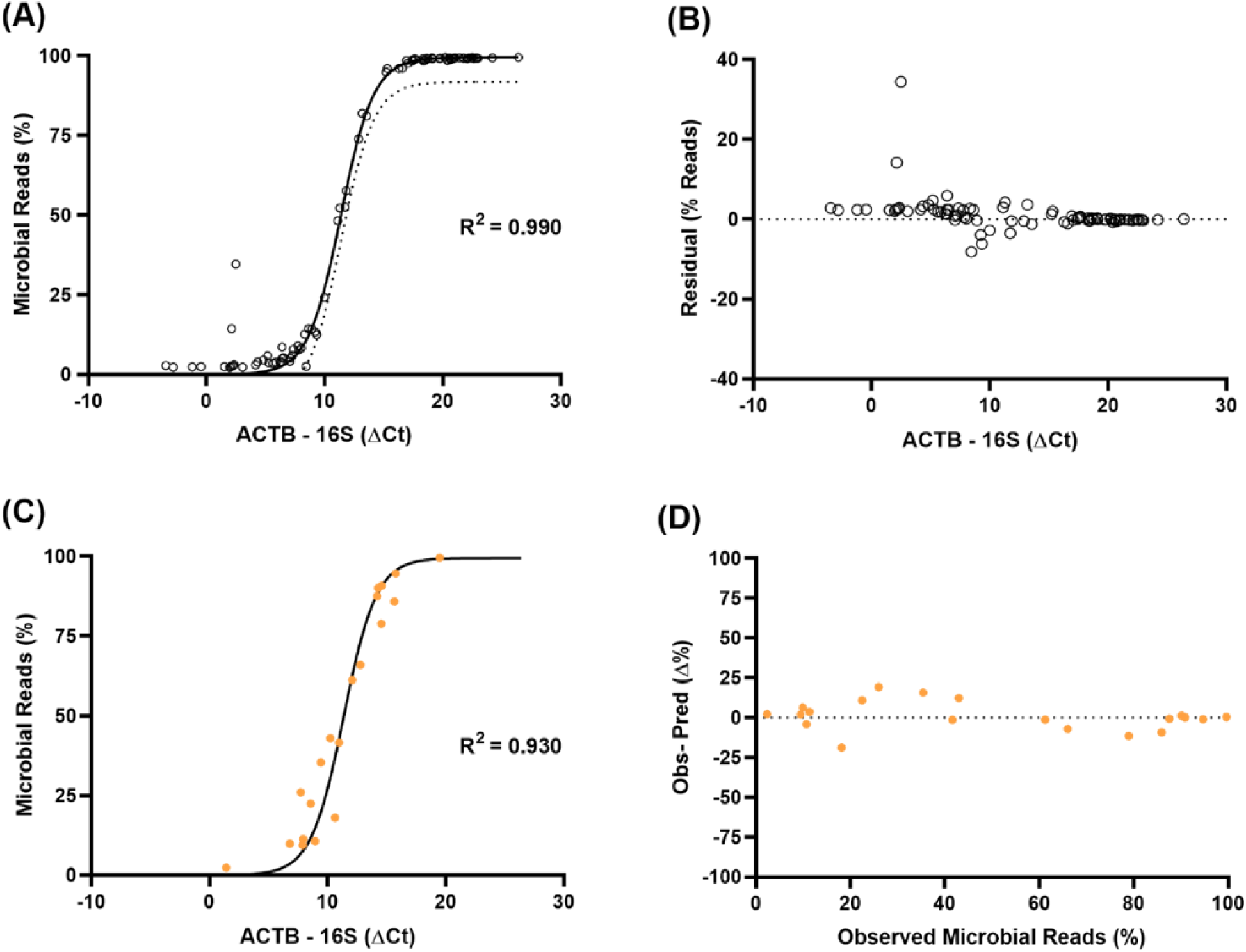
Statistical model to predict sample composition using qPCR prior to high-throughput sequencing. **(A)** Sigmoidal model generated from oropharyngeal swabs and stool samples depicting the relationship between the difference of human (ACTB) and bacterial (16S) qPCR values (Ct) with the percentage of microbial reads (R^2^ =0.990). Nonlinear regression line (solid) is based on the following logistic growth equation: % microbial reads = (2.7201549)/((99.50267)*e^(−0.7218*(ACTB-16S))+ 0.02733). One-tailed 95% prediction interval is depicted with a dotted line. **(B)** Model residuals. **(C)** Fitting of validation sample set on prediction model. The orange dots represent values derived from a validation sample set composed of vaginal secretions and rectal swabs samples and correlate well (R^2^ = 0.930) with the prediction model (solid black line). **(D)** Difference between expected and observed composition across the range of microbial content.

Using the equation derived from model E, we evaluated our approach on two different, independently collected sample types including vaginal secretions and rectal swabs. In **Fig 2C**, we show the relation between observed microbial reads percentages and the difference in Ct between 16S and ACTB qPCR, derived from our validation dataset, alongside a curve of expected values derived from model E. We observed the difference between predicted and observed microbial reads percentages to range from - 18.80% to +19.22% with a mean of +0.944% (**Supplementary figure 1B**). In **Fig 2D**, we show that this difference is consistent across the range of observed % microbial reads. Compared to the other models, model E best described the validation dataset, with a median difference of 0.25% and a standard deviation of 9.10% **(Supplementary table 1B)**. For comparison, model E described the initial sample set of oropharyngeal/stool samples with a median difference of 0.14% and a standard deviation of 4.35% **(Supplementary table 1A)**. Since the model performed similarly between the two datasets, we concluded that the model was able to describe a relation between 16S and β-actin qPCR and shotgun sequencing metagenomic data in a sample type-independent manner for microbial densities between 4% and 98%.We then developed a tool based on our model and the rarefaction curves on different samples type which predicts % microbial reads based on qPCR data and suggests a target number of reads based on sample type and desired analysis **(Supplementary)**.

The limitations of our study are as follows: The samples used in our study were low in fungal content. Therefore, our model may not accurately predict microbial content in sample sets where the majority of samples are rich in fungal content.

Moreover, as our results are based on protocols using specific reagents and technologies for both sequencing and qPCR, our tool may not accurately predict sequencing results when protocols, reagents, and/or technologies differ. However, given that we have established a robust link among 16S qPCR, B-actin qPCR, and sample content by sequencing, our approach can be easily adapted to fit different experimental settings.

## Conclusion

We have shown that shallowing shotgun sequencing depth can reduce measured alpha diversity in all measured sample types, with more diverse communities being more strongly negatively affected. We found that qPCR can function as a predictive tool for sample composition that was strongly correlated with shotgun sequencing data. We were able to create a model that can describe and predict variable sample types. We hope that our tool and methodology may help fellow researchers screen for sequenceable samples or allow for better optimization of sequencing.

## Methods

### qPCR

Samples were probed separately for the 16S rRNA gene, the 18S rRNA gene, and the human β-actin gene. All reactions were conducted in duplicate and RNase-free water was used as negative control. Each well contained 2 µL of sample DNA, 5 µL of Taqman Universal PCR mix (Applied Biosystems, Foster City, CA), 0.3 µM of forward primer, 0.3 µM of reverse primer, and 0.2 µM of primer probe. PCR was performed on a QuantStudio 6 Flex (Thermo Fisher Scientific, Waltham, MA) platform. Cycling was done as follows: 10 minutes at 95°C followed by 45 cycles of 95°C for 15 seconds and 60°C for 1 minute.

For 16S qPCR, we used forward primer “TCCTACGGGAGGCAGCAGT” (Invitrogen, Carlsbad, CA) and reverse primer “GGACTACCAGGGTATCTAATCCTGTT” (Invitrogen, Carlsbad, CA).(3) We used a FAM probe “CGTATTACCGCGGCTGCTGGCAC” with NFQ-MGB quencher (Applied Biosystems, Foster City, CA).(3)

For 18S qPCR, we used forward primer “GGRAAACTCACCAGGTCCAG” (Integrated DNA Technologies, Coralville, IA) and reverse primer “GSWCTATCCCCAKCACGA” (Integrated DNA Technologies, Coralville, IA).(1) We used a FAM probe “TGGTGCATGGCCGTT” with NFQ-MGB quencher (Applied Biosystems, Foster City, CA).(7) For human qPCR, we used a β-actin gene specific forward primer “CGGCCTTGGAGTGTGTATTAAGTA” (Invitrogen, Carlsbad, CA) and reverse primer “TGCAAAGAACACGGCTAAGTGT” (Invitrogen, Carlsbad, CA).(5) We used a VIC probe “TCTGAACAGACTCCCCATCCCAAGACC” with 3QSY quencher (Applied Biosystems, Foster City, CA).(8)

### Library preparation and sequencing

Libraries were prepared using Nextera Flex (Illumina, San Diego, CA) kits with the Nextera XT indices (Illumina, San Diego, CA). Barcoded sample libraries were pooled together to a concentration of 17.6 ng/ul which measured with a high-sensitivity DNA assay on a Qubit (Thermo Fisher Scientific, Waltham, MA) platform. A Mid-output reagent kit (Illumina, San Diego, CA) was used to sequence on the Miniseq, while a SP reagent kit (Illumina, San Diego, CA) was used on the Novaseq platform, both in 2×150bp mode.

### Read filtering and Taxonomic profiling

We filtered human reads from non-human reads using KneadData based on a human genome index for Bowtie 2 (9, 10). We considered sequence reads that did not match the database as microbial reads in our analyses. Taxonomic annotation was conducted using MetaPhlAn 2.0 and the ChocoPhlAn database (11). Rarefactions were performed using seqtk-1.3 to subsample the microbial reads of individual samples (12). Subsample compositions will be identified using MetaPhlAn2, and OTU tables were generated (11). Diversity indexes were calculated using Past 4 (13).

### Model generation

We used XLSTAT version 2019.4.2 (Addinsoft Inc., New York, NY) to generate multivariate linear regressions using either 16S and ACTB qPCR cycle and microbial reads percentages (Models A and C) or 16S, 18S, and ACTB qPCR cycle thresholds and microbial reads percentages (Models B and D). Multivariate linear regressions (models C and D) were also performed following a logit transformation of microbial reads percentages. Finally, for model E, we generated the non-linear regression model using the logistic growth equation in GraphPad Prism version 8.3.0 for Windows (GraphPad Software, San Diego, CA).

## Supporting information

Supplementary Tool

---

**Supplementary Figure 1. (1A)** Residuals for 5 multivariate models generated using a sample set comprised of oropharyngeal swabs and stool samples. i) Model A represents a linear fit taking into account microbial and human-derived qPCR values; ii) model B represents a linear fit taking into account microbial, fungal, and human-derived qPCR values; iii) model C represents a linear fit taking into account microbial and human-derived qPCR values after a logit transform of the data; iv) model D represents a linear fit taking into account microbial, fungal, and human-derived qPCR values after a logit transform of the data; and v) model E represents a nonlinear regression model based on the logistic growth equation taking into account microbial and human-derived qPCR values. Error bars depict 1 standard deviation centered around the mean. **(1B)** Difference between observed and predicted percentage of microbial reads, by model, using a validation dataset comprised of independently collected rectal swabs and vaginal secretion samples.

**Supplementary Table 1. (1A)** Table summarizing residual values and model type of the 5 statistical models (A-E) tested in this study. **(1B)** Residual values of the 5 models when applied to an independent, validation dataset.

