## Supplementary Figures and Table for "Two-target quantitative PCR to predict library composition for shallow shotgun sequencing"

### Slide 1
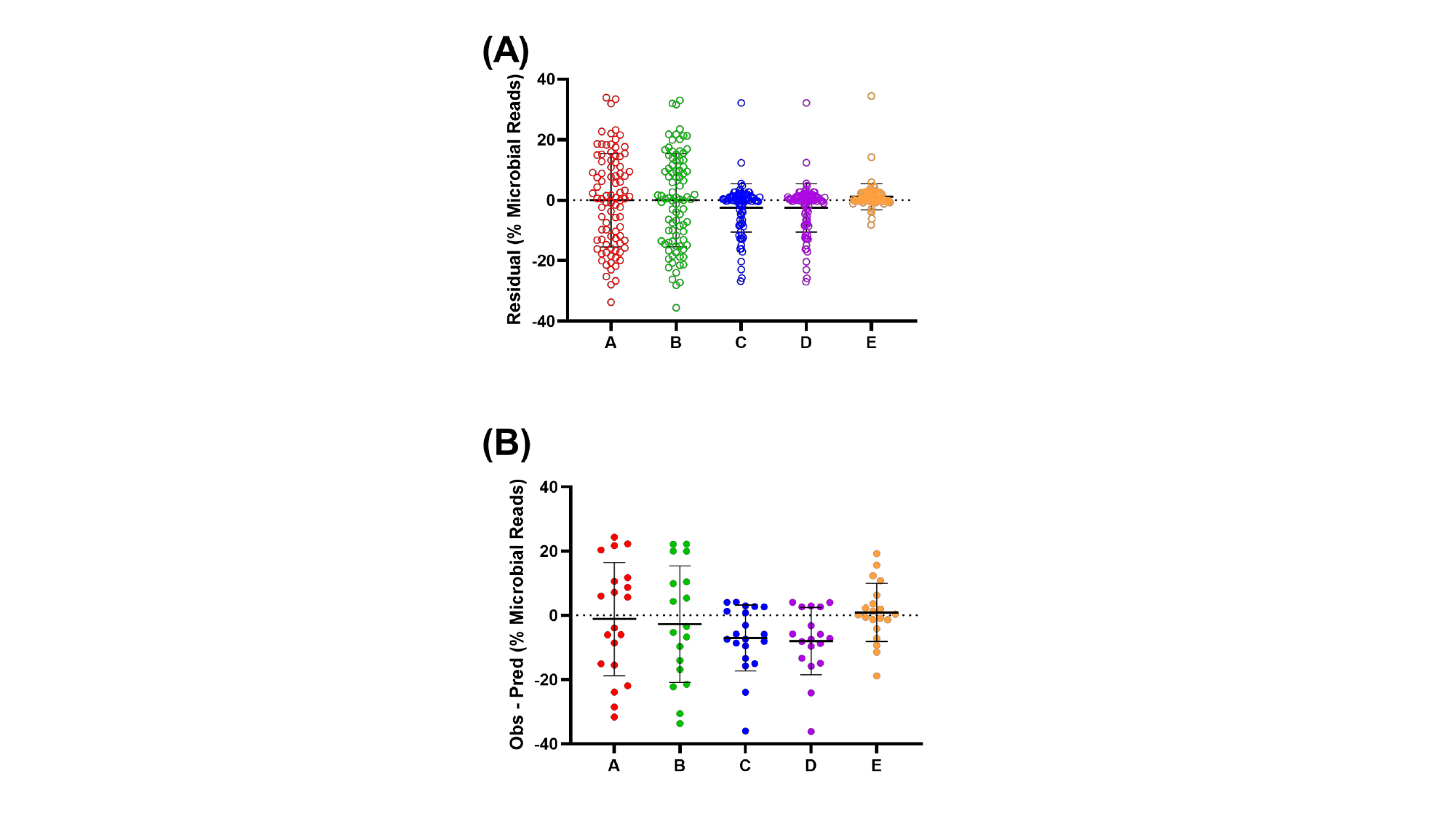

### Slide 2
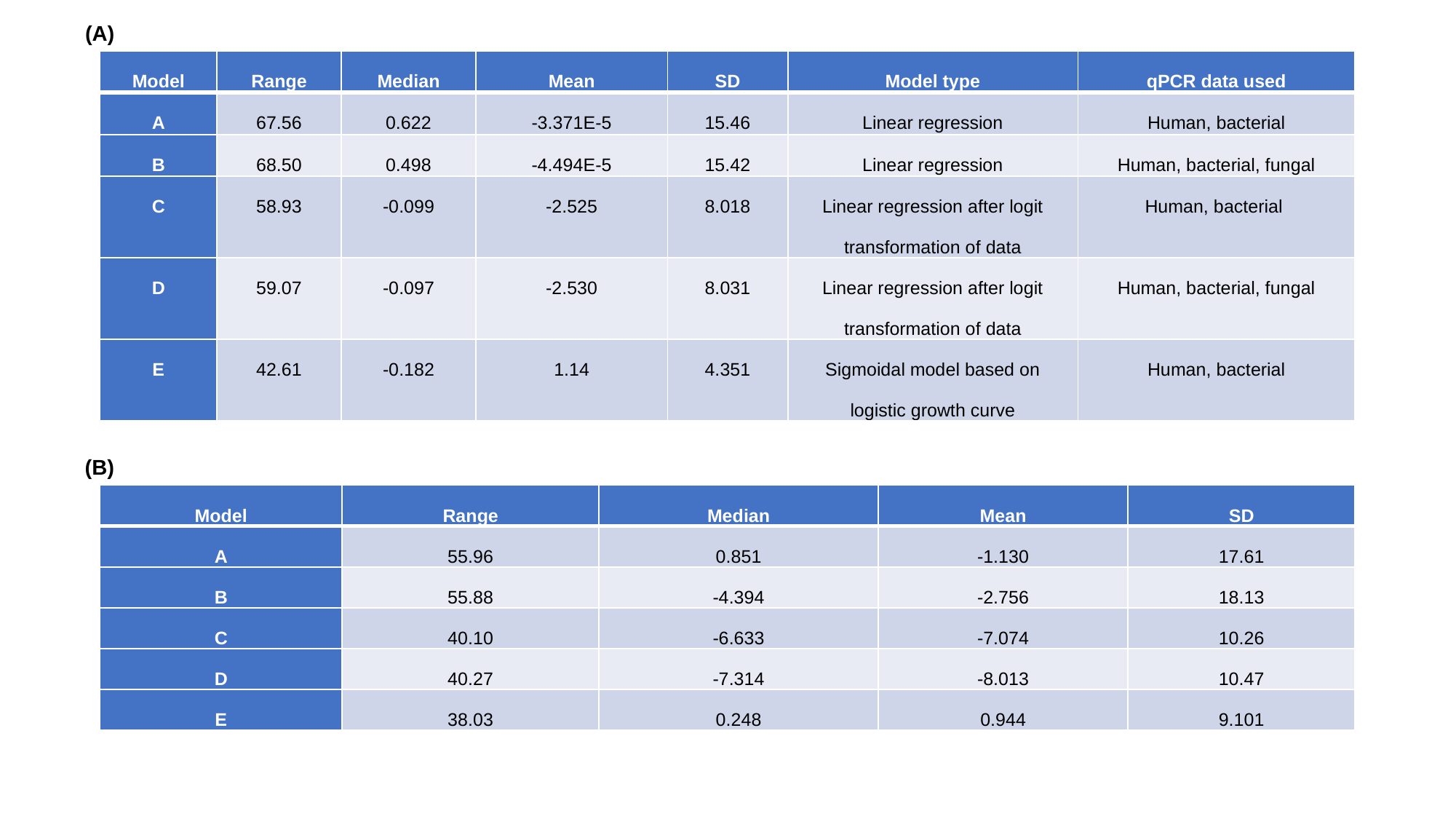

(A)
| Model | Range | Median | Mean | SD | Model type | qPCR data used |
| --- | --- | --- | --- | --- | --- | --- |
| A | 67.56 | 0.622 | -3.371E-5 | 15.46 | Linear regression | Human, bacterial |
| B | 68.50 | 0.498 | -4.494E-5 | 15.42 | Linear regression | Human, bacterial, fungal |
| C | 58.93 | -0.099 | -2.525 | 8.018 | Linear regression after logit transformation of data | Human, bacterial |
| D | 59.07 | -0.097 | -2.530 | 8.031 | Linear regression after logit transformation of data | Human, bacterial, fungal |
| E | 42.61 | -0.182 | 1.14 | 4.351 | Sigmoidal model based on logistic growth curve | Human, bacterial |
(B)
| Model | Range | Median | Mean | SD |
| --- | --- | --- | --- | --- |
| A | 55.96 | 0.851 | -1.130 | 17.61 |
| B | 55.88 | -4.394 | -2.756 | 18.13 |
| C | 40.10 | -6.633 | -7.074 | 10.26 |
| D | 40.27 | -7.314 | -8.013 | 10.47 |
| E | 38.03 | 0.248 | 0.944 | 9.101 |
